# Hematologic and systemic metabolic alterations due to Mediterranean type II G6PD deficiency in mice

**DOI:** 10.1101/2021.05.31.446353

**Authors:** Angelo D’Alessandro, Heather L Howie, Ariel M. Hay, Karolina H. Dziewulska, Benjamin Brown, Matthew J Wither, Matthew Karafin, Elizabeth F. Stone, Steven L Spitalnik, Eldad A Hod, Richard O Francis, Xiaoyun Fu, Tiffany Thomas, James C Zimring

## Abstract

Deficiency of Glucose 6 phosphate dehydrogenase (G6PD) is the single most common enzymopathy, present in approximately 400 million humans (e.g. 5% of humans). Its prevalence is hypothesized to be due to conferring resistance to malaria. However, G6PD deficiency also results in hemolytic sequelae from oxidant stress. Moreover, G6PD deficiency is associated with kidney disease, diabetes, pulmonary hypertension, immunological defects, and neurodegenerative diseases. To date, the only available mouse models have decreased levels of G6PD due to promoter mutations, but with stable G6PD. However, human G6PD mutations are missense mutations that result in decreased enzymatic stability. As such, this results in very low activity in red blood cells and platelets that cannot synthesize new protein. To generate a more accurate model, the human sequence for a severe form of G6PD deficiency (Med -) was knocked into the murine G6PD locus. As predicted, G6PD levels were extremely low in RBCs and deficient mice have increased hemolytic sequalae to oxidant stress. G6PD levels were mildly decreased in non-erythroid organs, consistent with what has been observed in humans. Juxtaposition of G6PD deficient and wild-type mice revealed altered lipid metabolism in multiple organ systems. Together, these findings both establish a new mouse model of G6PD deficiency that more accurately reflects human G6PD deficiency and also advance our basic understanding of altered metabolism in this setting.

## Introduction

Glucose-6-phosphate dehydrogenase (G6PD) is the first and rate determining enzyme in the pentose phosphate pathway (PPP), which utilizes glucose to generate NADPH; the latter is the major reducing equivalent that fuels multiple pathways by which cells handle oxidative stress. Deficiency in the activity of G6PD is the single most common enzymopathy in humans, estimated to be present in approximately 400 million people worldwide(1). The complete absence of G6PD is not compatible with life, and the vast majority of mutations leading to G6PD deficiency in humans are missense mutations leading to an unstable G6PD enzyme. Based upon the mutation and resulting G6PD levels, disease can range from mild to severe deficiency (2, 3). RBCs from humans with G6PD deficiency are particularly susceptible to oxidant stress for two prevailing reasons. First, because RBCs lack mitochondria, the PPP is their main source of NADPH. Second, mature RBCs are unable to synthesize new proteins. When G6PD deficient humans encounter an illness or consume a drug or food that increases oxidant stress (e.g., antimalarial quinone drugs or fava beans), they can manifest symptoms of actue hemolysis, ranging from mild to lethal(4, 5). Recent findings have also linked G6PD deficiency to many other diseases outside the RBC compartment, including immunological(1), cardiovascular(6), endocrine(7), renal(7), neurological(8), and pulmonary pathologies(9).

Because G6PD deficiency is so prevalent in humans, and because its biology remains poorly understood, a translatable animal model of G6PD deficiency is essential. Two separate mouse models of G6PD deficiency have been described(10, 11), both derived by random mutagenesis. Importantly, both result in altered levels of G6PD expression, but with a normal amino acid sequence and normal protein stability(10, 11). As such, they do not recapitulate the human situation where young RBCs have high levels of G6PD activity, which then diminish as the RBCs age. To our knowledge, no mouse model has been described with an unstable G6PD enzyme.

Type II G6PD deficiency is the most severe form that lacks ongoing non-spherocytic hemolytic anemia at baseline. The best studied type II deficiency is the “Mediterranean variant” that results from a Ser 188 to Phe mutation (12, 13). Herein, we describe a novel mouse in which the genomic sequence for the human “Mediterranean-Type” G6PD-deficient mutant was knocked into the murine G6PD locus, thus recapitulating the human enzymopathy in a murine system. The phenotype of this mouse resembles that of humans with G6PD deficiency, in that the mice are healthy at baseline but demonstrate hemolysis upon challenge with oxidant stress. A detailed analysis of these mice was performed, under normal and stressed conditions, including metabolomics profiling of multiple organs. In addition to describing a new model, these findings provide novel understanding of the metabolic consequences of G6PD deficiency in multiple organ systems, as well as unique insights into the biochemical mechanisms by which oxidant stress alters RBCs, both in the G6PD-normal and G6PD-deficient states.

## Results

### Generation of a Novel Murine Model of Human G6PD Deficiency

A targeting construct was generated to insert the human type II Mediterranean variant [hMed(-)] variant directly into the murine G6PD locus (**Figure 1A**.**1**). To maintain genetic structure, the entire genomic sequence of human hMed(-) was utilized from exons 3-12. Out of concern for disrupting genomic regulatory elements in the proximal murine sequence, exons 1 and 2, and introns 1 and 2 of the murine sequence were left unaltered. As such, the final G6PD gene product is the hMed(-) form (Ser188Phe) that also has two amino acids in the N-terminus from the murine sequence (**Figure 1B**).

**Figure 1.**
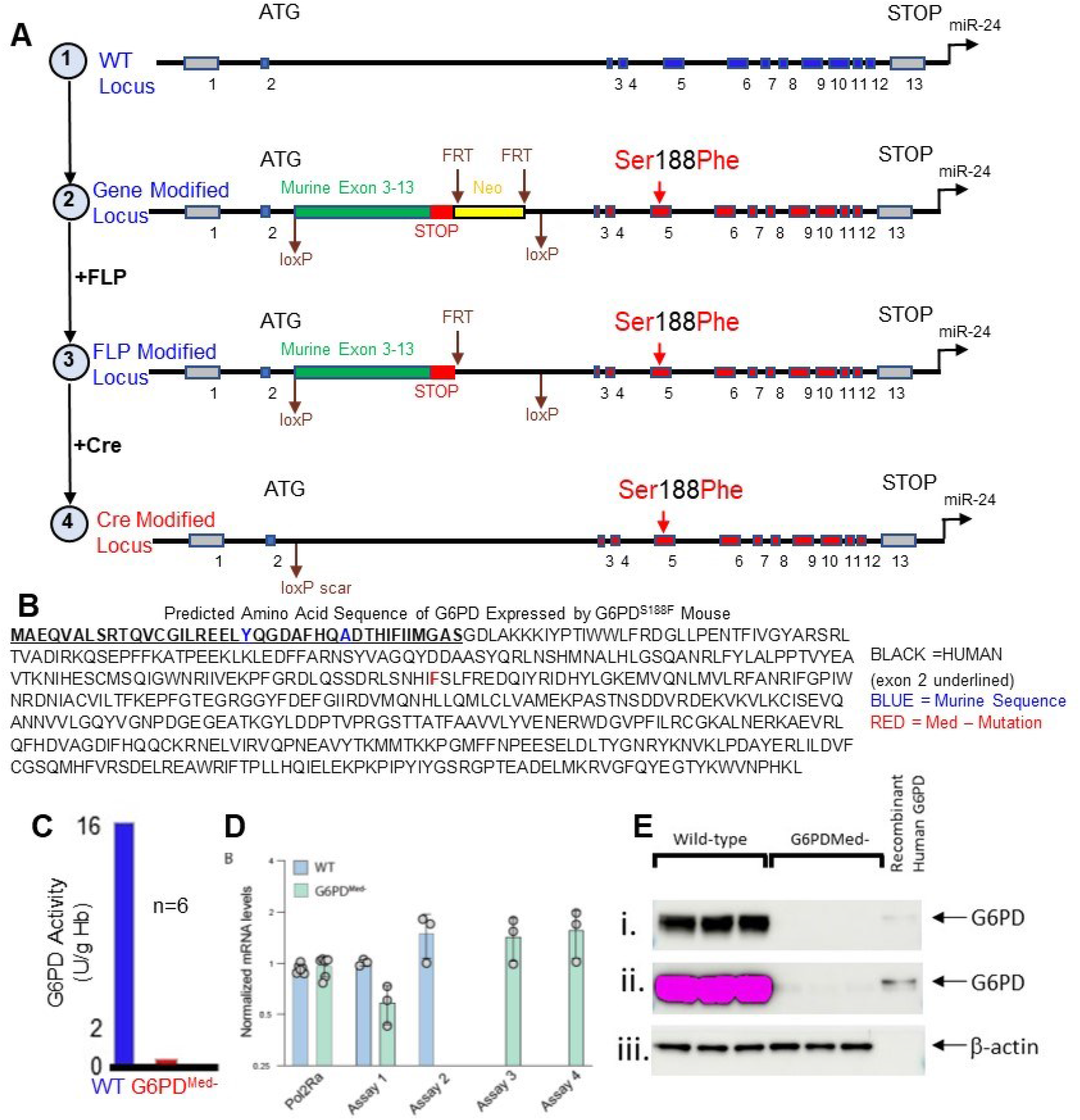
Generation of a new G6PD deficient mouse model. **(A)** Schematic representation of the WT G6PD locus (top) and related modifications (bottom). **(B)** Predicted protein sequence of knocked in G6PD gene. **(C)** G6PD activity in wild-type vs knockin mouse. **(D)** mouse vs. human mRNA levles in G6PD_Med-_ and WT mice, **(E)** Western blot analysis of cytoplasm from RBCs. Because the G6PD gene is X lined, males were used in all cases. G6PD_Med-_ mice were hemizygous for the knocked in human gene, WT mice are litter mate contols with the mouse G6PD.

Out of concern that the modification may be embryonic lethal, and also to allow additional experimental flexibility, the genetic construct was designed for conditional expression of the hMed(-) gene. Wild-type mouse G6PD cDNA (exons 3-13) were inserted upstream of the human gene sequence and were flanked with LoxP sites. This construct is designed to express wild-type murine G6PD until it is exposed to CRE recombinase, at which time the mouse cDNA is excised and the human genomic sequence for the hMed(-) G6PD is expressed. (**Figure 1A**.**3-1A**.**4**). This mouse is called “Con-hMed(-)”, and was generated using ES cells from a C57BL/6 background. Southern blot analysis was carried out both on modificed ES cells and also on founder mice to confirm the genetic alteration was made (**Supplementary Figure 1A**).

The Con-hMed(-) mouse was bred with mice that expressed germline CRE, and then were crossed with wild-type B6 to remove the CRE gene (**Figure 1A**.**4**), resulting in a germline hMed(-) mouse (called G6PD_Med-_) . Correct excision of the floxed region was confirmed by PCR (data not shown). Male and female G6PD_Med-_ mice were viable and fertile, and females had normal fecundity. As in humans, the murine G6PD locus is on the X-chromosome; thus, heterozygous female G6PD_Med-_ mice were bred with wild-type males, such that 50% of the male offspring were hemizygous for hMed(-) and the other 50% were wild-type. In this way, G6PD_Med-_ mice were compared to littermate control wild-type mice.

RBCs from G6PD_Med-_ mice had only 5% of the G6PD activity compared to wild-type controls (**Figure 1C**). The decreased G6PD activity was not due to decreased gene expression, as there was no difference in the amount of hMed(-) G6PD mRNA in G6PD_Med-_ compared to wild-type G6PD mRNA in B6 mice (Figure 1D). However, Western blot analysis of the cytoplasm from RBCs using an antibody that reacts with both human and murine G6PD demonstrated only trace G6PD in RBCs from G6PD_Med-_ mice (Figure 1E.i) even intentional over expression (Figure 1E.ii.) This was not due to differences in loading the gel, as equivalent levels of beta actin were detected in all samples.

Peripheral blood was analyzed for standard metrics of RBC biology. There was no statistical differences in the values of RBC number, Hemoglobin concentration, hematocrit, mean corposcular volume, red cell distribution width, or reticulocyte counts between G6PD_Med-_ and wild-type mice (**Supplementary figure 2**).

**Figure 2.**
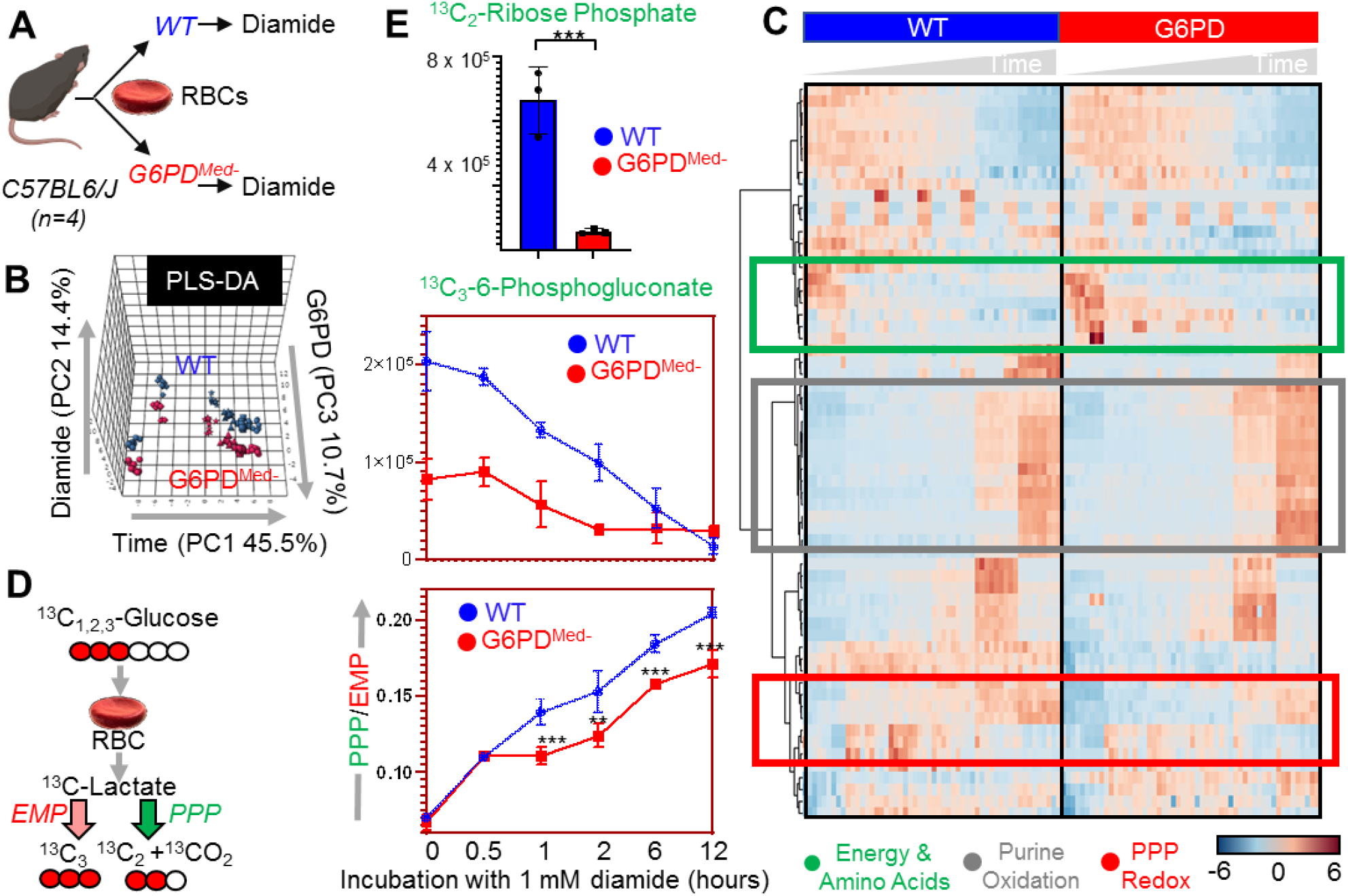
Metabolic effect of diamide challenge in RBCs from WT and G6PD deficient mice. RBCs from WT and G6PD def mice were incubated with 1mM diamide (**A**). RBCs were tested at 0min (no diamide), 30min, 1h, 2h, 6h and 12h from incubation with diamide prior to metabolomics analysis. Multivariate analyses included PLS-DA (**B**) and hierarchical clustering analysis (**C**) clearly indicate a time-dependent effect of the treatment on RBCs (PC1 explaining 45.5% of the total variance) and highlighted the impact of G6PD activity (PC3 – 10.7% of the total variance). Significant metabolites by repeated measures one-way ANOVA are shown in the heat map in **C**. In **D**, the experiment was repeated by incubating RBCs in presence of 1,2,3-^13^C_3_-glucose. By quantifying isotopologues M+2 and M+3 of lactate (and the relative ratio), fluxes through Pentose Phosphate Pathway vs Glycolysis can be determined, confirmed a significantly lower activation of this pathway in RBCs from G6PD def mice upon diamide challenge.

### -hMed(-) RBCs have decreased PPP activity ex vivo

To test how decreased G6PD activity in G6PD_Med-_ RBCs affected the PPP, RBCs from wild-type and G6PD_Med-_ RBCs were exposed to an oxidant challenge (1 mM diamide) for 12 hours in the presence of [1,2,3]-^13^C_3_-glucose (**Figure 2A**). Multivariate analyses, including partial least square-discriminant analysis (PLS-DA – **Figure 2B**) and hierarchical clustering analysis (**Figure 2C**), showed significantly distinct metabolic phenotypes between G6PD_Med-_ RBCs and wild-type RBCs. A version of the heat map in **Figure 2C**, which also includes the top 50 significant metabolites (and isotopologues) that are significant by repeat measure ANOVA, is provided as **Supplementary Figure 3**. Results are also provided in tabular form in **Supplementary Table 1**. G6PD_Med-_ mice had a substantially decreased ratio of ^13^C_2_/^13^C_3_ lactate isotopologues, indicating decreased flux through the PPP (**Figure 2**.**D**). Consistent with this interpretiaion of the decreased ratio of ^13^C_2_/^13^C_3_ lactate isotopologues, RBCs from G6PD_Med-_ mice also had significantly lower ribose phosphate **(Figure 2E)**, which is an end product of the PPP, as well as ^13^C_3_-phosphogluconate, which is an intermdiate of the oxidative phase of the PPP. The untargeted metabolomics analyses also demonstrate that, in addition to alterations in glycolysis and PPP, hMed(-) RBCs had significantly altered redox regulated pathways, including methionine metabolism and purine oxidation (**Figure 3**). As these are steady state levels, one cannot infer changes in sythensis or consumption, only that levels are different.

**Figure 3.**
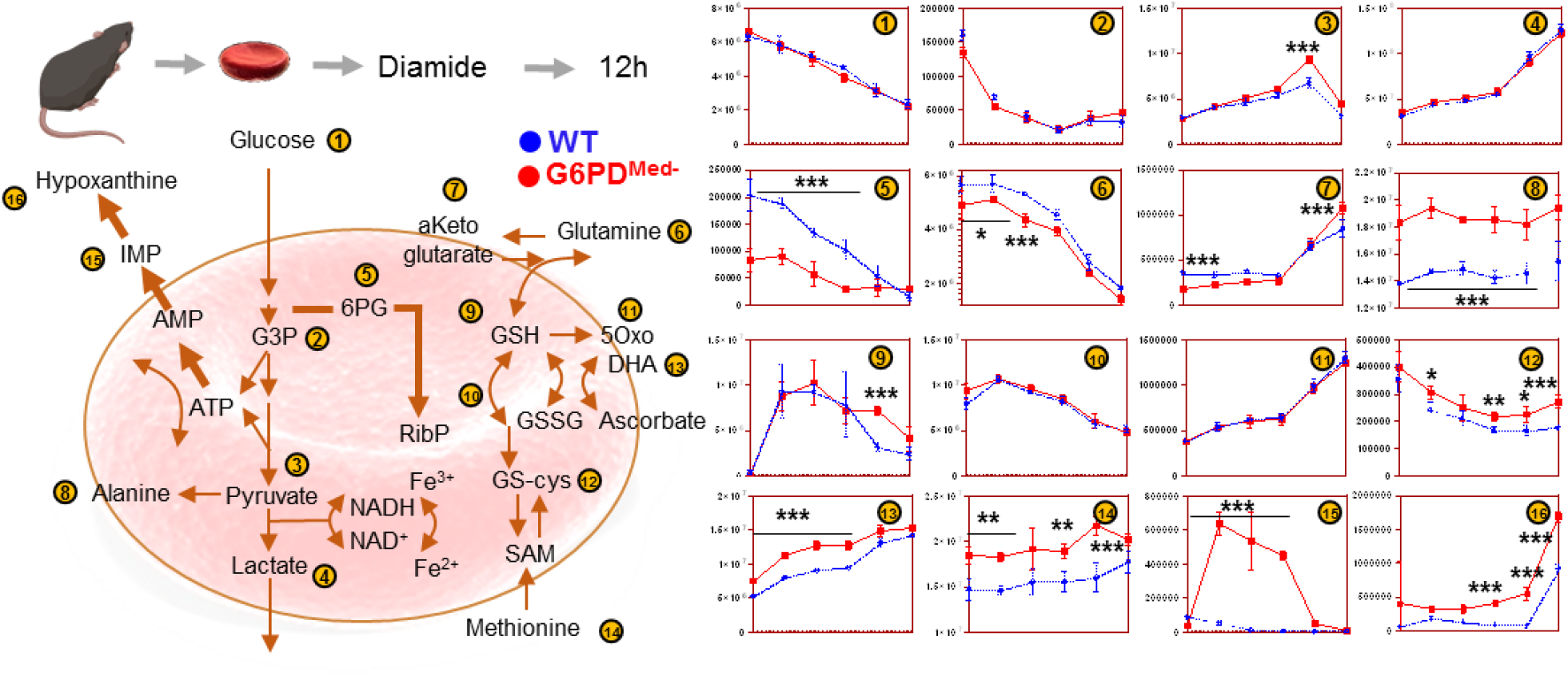
Overview of glycolysis, pentose phosphate pathway, glutathione metabolism and recycling, purine oxidation in RBCs from WT (blue) and G6PD def (red) mouse red blood cells (n=3) treated with 1 mM diamide for up to 12 hours. Time points tested were 0 (no diamide), 30min, 1h, 2h, 6h and 12h from incubation with diamide. Significantly increased pyruvate/lactate ratios and alanine accumulation in G6PD def mouse RBCs is consistent with an increased utilization of NADH by other enzymes than lactate dehydrogenase, consistent with higher levels/activity of methemoglobin reductase in prior human studies on Med-G6PD def RBCs (Tzounakas et al. 2016 - https://www.ncbi.nlm.nih.gov/pubmed/27094493).

### RBCs from G6PD_Med-_ mice have increased hemolysis in response in vivo to pharmacological oxidant stress

To test the hypothesis that G6PD_Med-_ mice were more sensitive to oxidant stress *in vivo*, a method was used by which the complete blood compartment of G6PD_Med-_ mice and wild-type wer biotinylated (14). This is the equivalent of an in vivo pulse chase experiment, allowing the determination of RBC lifespan without interference of erythropoiesis, as newly generated RBCs are not biotinylated. Baseline studies determined that the normal circulatory lifespan of G6PD_Med-_ RBCs was indistinguishable from that of wild-type controls (**Figure 4A**).

**Figure 4.**
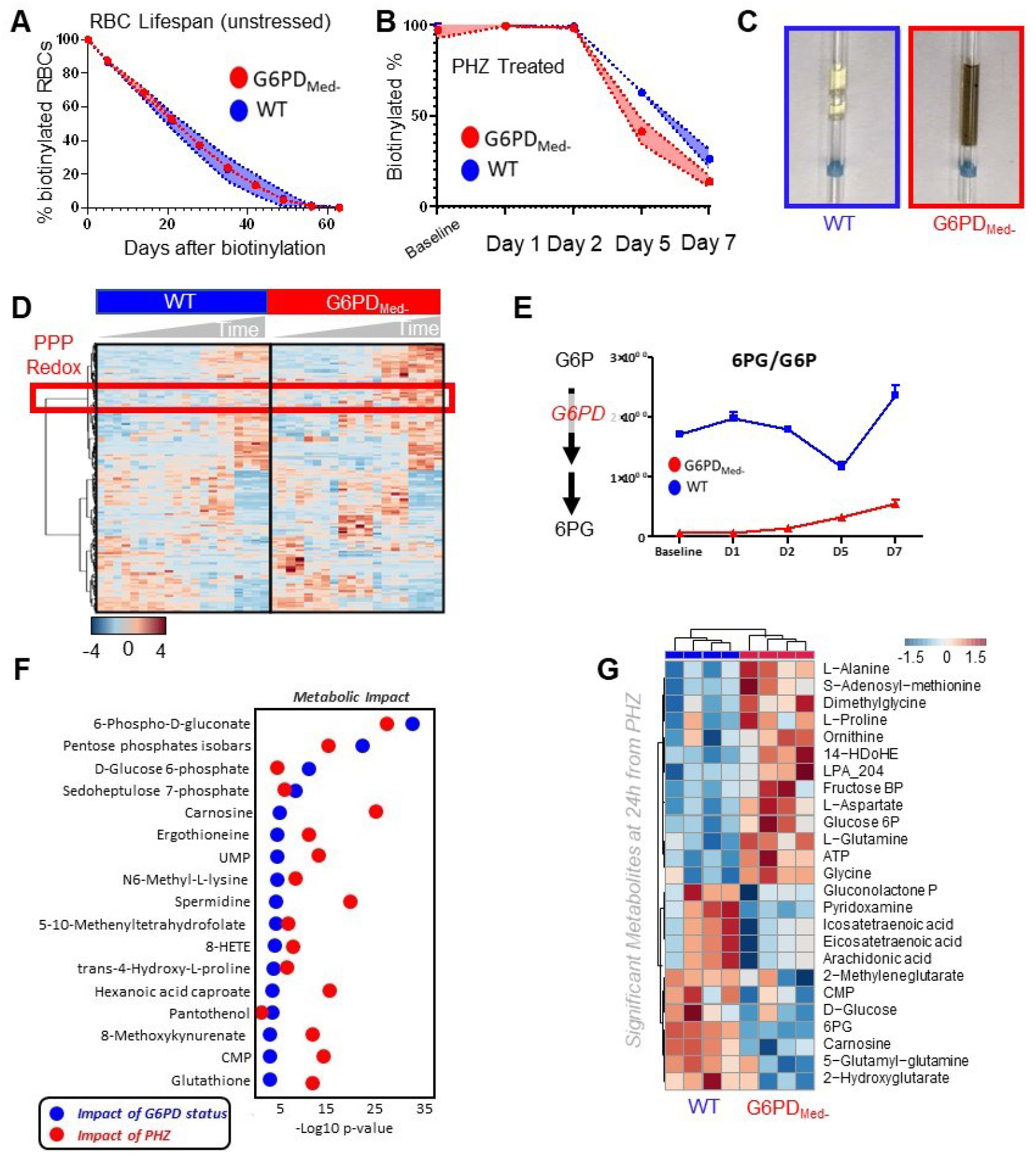
Phenylhydrazine (PHZ) induces brisk hemoglobinuria in G6PD def mouse RBCs, but not in WT. **(A).** Biotinylation experimetns showed significant decreases in circulating RBC survival after the 2^nd^ PHZ treatment in G6PD deficient mice (B). G6PD def RBCs at baseline and upon treatment with PHZ for up to 7 days are incapable of activating the PPP (C), as determined by the ratios of G6PD product/substrate, a proxy for the determination of the enzyme activity by law of mass action, as described (Zhang et al. Sci Signaling 2018 - https://www.ncbi.nlm.nih.gov/pubmed/29991649) (D). Significant metabolic changes were noted between the two groups upon prolonged treatment with PHZ (C). Pathway analysis highlighted 4 PPP-related metabolites on the top 5 of significant metabolic changes by two way ANOVA (first column: impact of G6PD deficiency; second column: impact of PHZ treatment; third column: interactions: (E-F)). These changes mostly involved oxidant stress and sulfur metabolism-related pathways (ergothioneine, glutathione, oxylipins) and ultimately resulted in distinct metabolic phenotypes as a function of PHZ treatment and between WT and G6PD def RBCs. These differences are clear when comparing the time point at 24h from PHZ injection in WT and G6PD def mice as highlighted in the heat map in F.

Phenylhydrazine (PHZ) is a classic chemical to induce oxidant stress in vivo, and is known to cause intravascular hemolysis in human G6PD-deficient subjects(15). PHZ induced a significantly faster rate of RBC clearance from circulation of G6PD_Med-_ mice compared to wild-type controls (**Figure 4B – Supplementary Table 2**). In addition, G6PD_Med-_ but not wild-type controls displayed significant hemoglobinuria (**Figure 4C**). Metabolomics analyses were performed in blood from the mice at baseline (prior to biotinylation), at day 1 (after biotinylation), at day 2 (middle of PHZ challenge), and at days 5 and 7 (post-PHZ challenge). Results are summarized in the heat map in **Figure 4D**, with more extensive information provided in tabulated and vectorial forms (including metabolite names) in **Supplementary Figure 4**.

The ratio of steady state glucose 6-phosphate (G6P) to 6-phospho-gluconate (6PG) (i.e., the substrate and downstream metabolite of G6PD, respectively), is a proxy for G6PD activity(16) (**Figure 4E**). Consistent with the *ex vivo* measurements described above, *in vivo* measurements of 6PG/G6P ratios were significantly lower at baseline and following PHZ exposure in G6PD_Med-_ mice, compared to wild-type animals (**Figure 4E**). Repeated measure ANOVA of time course data further highlighted that the top four metabolites that significantly differed between the two groups as a function of PHZ stimulation were PPP metabolites (i.e., G6P, 6PG, ribose phosphate and isobaric isomers, sedoheptulose phosphate; **Figure 4F**). Other metabolites related to redox homeostasis were significantly affected by PHZ stimulation, as a function of G6PD status, including glutathione, oxylipins (HETEs), proteolysis markers (e.g., hydroxyproline, methyl-lysine), and other antioxidant metabolites (e.g., carnosine, ergothioneine; **Figure 4E, Supplementary Figure 4**). These changes between G6PD_Med-_ and wild-type mice were most evident after PHZ injection (a heat map of the top 25 significant metabolites by unpaired t-test for this time point is provided in **Figure 4G**). A detailed representation of these metabolites and related pathways as a function of the entire time course is provided in the form of line plots in **Figure 5**.

**Figure 5.**
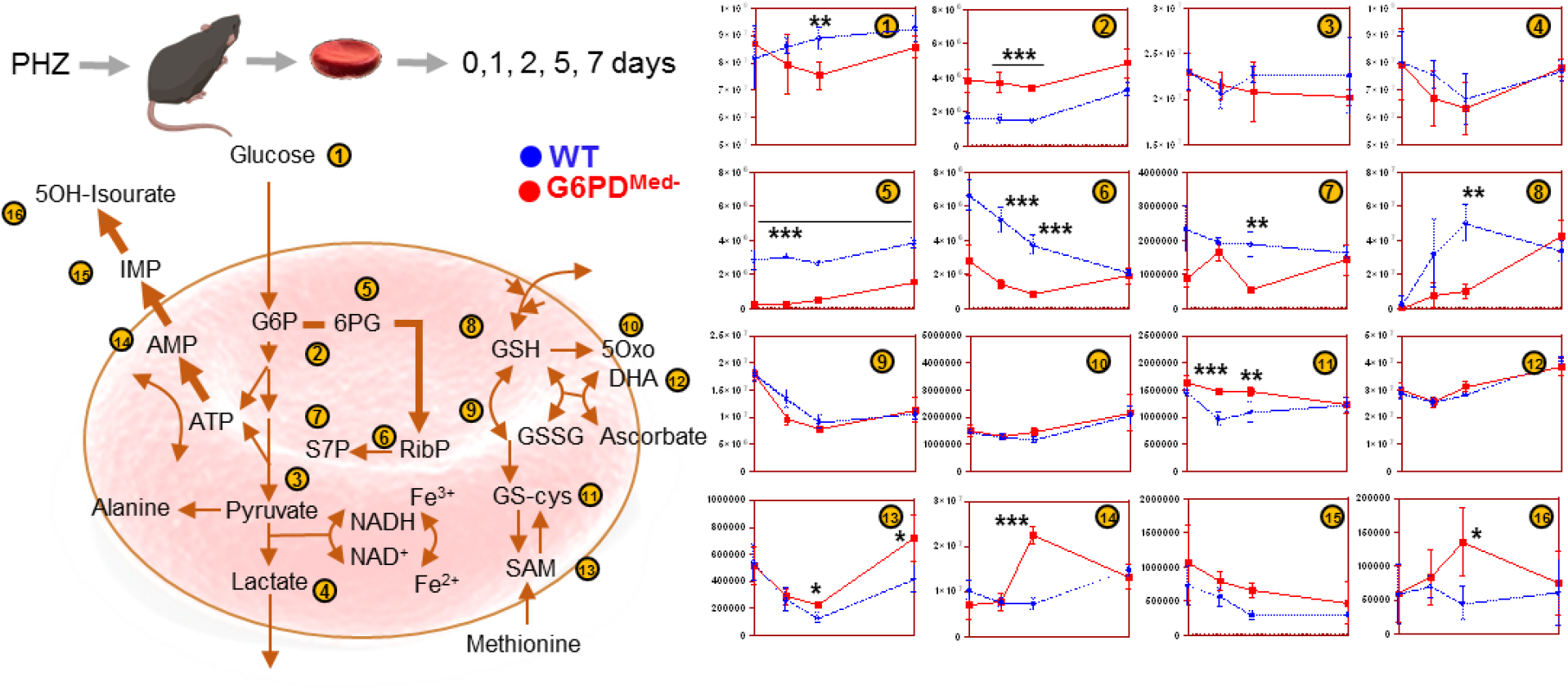
Metabolic impact of PHZ treatment in vivo on WT and G6PD def mouse RBCs. Overview of glycolysis, pentose phosphate pathway, glutathione metabolism and recycling, purine oxidation in RBCs from WT (blue) and G6PD def (red) mouse red blood cells (n=3) treated with 1 mM phenylhydrazine (PHZ) for up to 7 days. Time points tested were 0 (no PHZ), 1, 2, 5 and 7 days from treatment with PHZ.

### RBCs from G6PD_Med-_mice have normal post-transfusion recovery after storage

Refrigerated storage of human RBCs is an iatrogenic source of oxidant stress that is a necessary logistical component of blood banking and clinical transfusion practice. It was recently reported that when RBCs are stored from humans with G6PD deficiency and are then transfused, they have a modest (i.e., ∼6%), but statistically significant, decrease in 24-hour post-transfusion recovery. To test how the hMed(-) mutation affects storage of murine RBCs, donor mice were exsanguinated and RBCs were stored using an established methodology that models the human setting(17). In this case, the 24-hour post-transfusion recovery was the same for G6PD_Med-_ and wild-type mice (**Figure 6A**).

**Figure 6.**
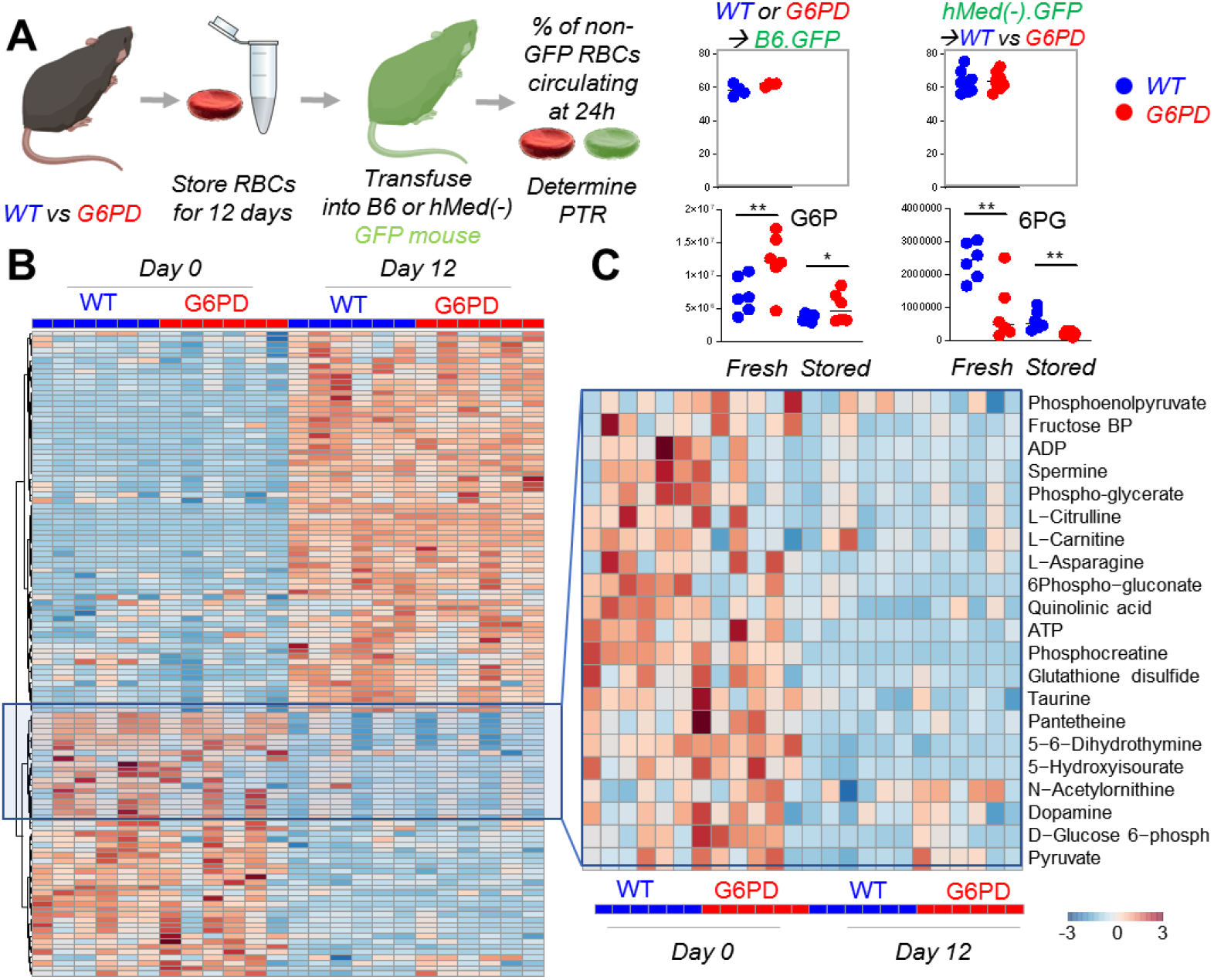
G6PD deficient mouse RBCs have comparable metabolism and post-transfusion recovery to wild-type RBCs at the end of storage. RBCs from WT and hMed(-) G6PD mice (n=6) were stored under conditions mimicking storage in the blood bank for 12 days (**A**). At the end of storage, RBCs were transfused into Ubi-GFP mice and flow cytometry studies were performed to determine the percentage of transfused RBCs circulating at 24h from transfusion, which was determined to be comparable between the two groups. Transfusing hMed(-).GFP into WT or hMed(-) non-GFP recipient did not impact the PTR % measurement. Metabolic phenotypes of G6PD deficient RBCs showed some significant differences at baseline (especially with respect to glycolysis, the pentose phosphate pathway and glutathione homeostasis (B). However, these changes were not appreciable by the end of storage. A detail for glucose 6-phosphate (G6P) and 6-phosphogluconate (6PG) is provided in (C)

The human studies that demonstrated decreased 24-hour recoveries of transfused RBCs from G6PD-deficient donors used autologous transfusions of radiolabeled RBCs; as such, the G6PD-deficient RBCs were not only exposed to the oxidative stress of storage, but were also introduced into a recipient with potentially altered redox biology due to G6PD-deficiency. To allow for autologous transfusion studies in mice, G6PD_Med-_ mice were crossed with B6.GFP mice. RBCs from G6PD_Med-_ mice were then collected, stored, and transfused into either B6.GFP mice with normal G6PD or into G6PD_Med-_.GFP recipients; in this approach, transfused RBCs are enumerated as the GFP-negative population. No difference in 24-hour post-transfusion recovery was observed in G6PD_Med-_.GFP, as compared to B6.GFP recipients, ruling out that differences in storage biology would be present if both donor and recipient were hMed(-) (**Figure 6A**).

Stored RBCs were also analyzed by metabolomics. At baseline (i.e., before storage), differences were observed in glycolysis and the PPP, glutathione homeostasis, and purine and amino acid metabolism (**Figure 6**.**B**); specifically, genotype-dependent alterations were detected with regards to arginine, tryptophan, and tyrosine metabolism, as gleaned by differential levels of citrulline, indoles, and dopamine, respectively (**Figure 6B** – right hand panel). Although increases in G6P and decreases in 6PG were significant in G6PD_Med-_ mice both at baseline and at the end of storage (12 days), at day 12 the fold-changes for both metabolites between the two genotypes were negligible (**Figure 6**.**C**).

### Effects of hMed(-) on non-erythroid tissues

G6PD deficiency has often been considered to be RBC specific, because, unlike RBCs that cannot synthesize protein, nucleated cells can compensate for decreased G6PD enzyme activity by increasing synthesis. However, in humans with G6PD deficiency, other tissues (e.g., muscle and endothelial cells in the pulmonary artery) have decreased G6PD activity, albeit not as severe as seen in RBCs(18, 19). To perform a detailed analysis of multiple organs, which is not feasible in human studies, metabolomics analyses were performed on brain, heart, kidney liver, and spleen of G6PD_Med-_ compared to wild-type mice (**Figure 7A**). Detailed results from hierarchical clustering analyses of all the metabolomics data are provided for each organ in **Supplementary Figures 5-10** and, in tabulated form, in **Supplementary Table 3** (significant metabolites in **Supplementary Table 4**). An overview of the multivariate analyses, including PLS-DA and heat maps reporting the top 25 significant metabolites by unpaired t-test, is provided for each organ in **Figure 7-B-F**.

**Figure 7.**
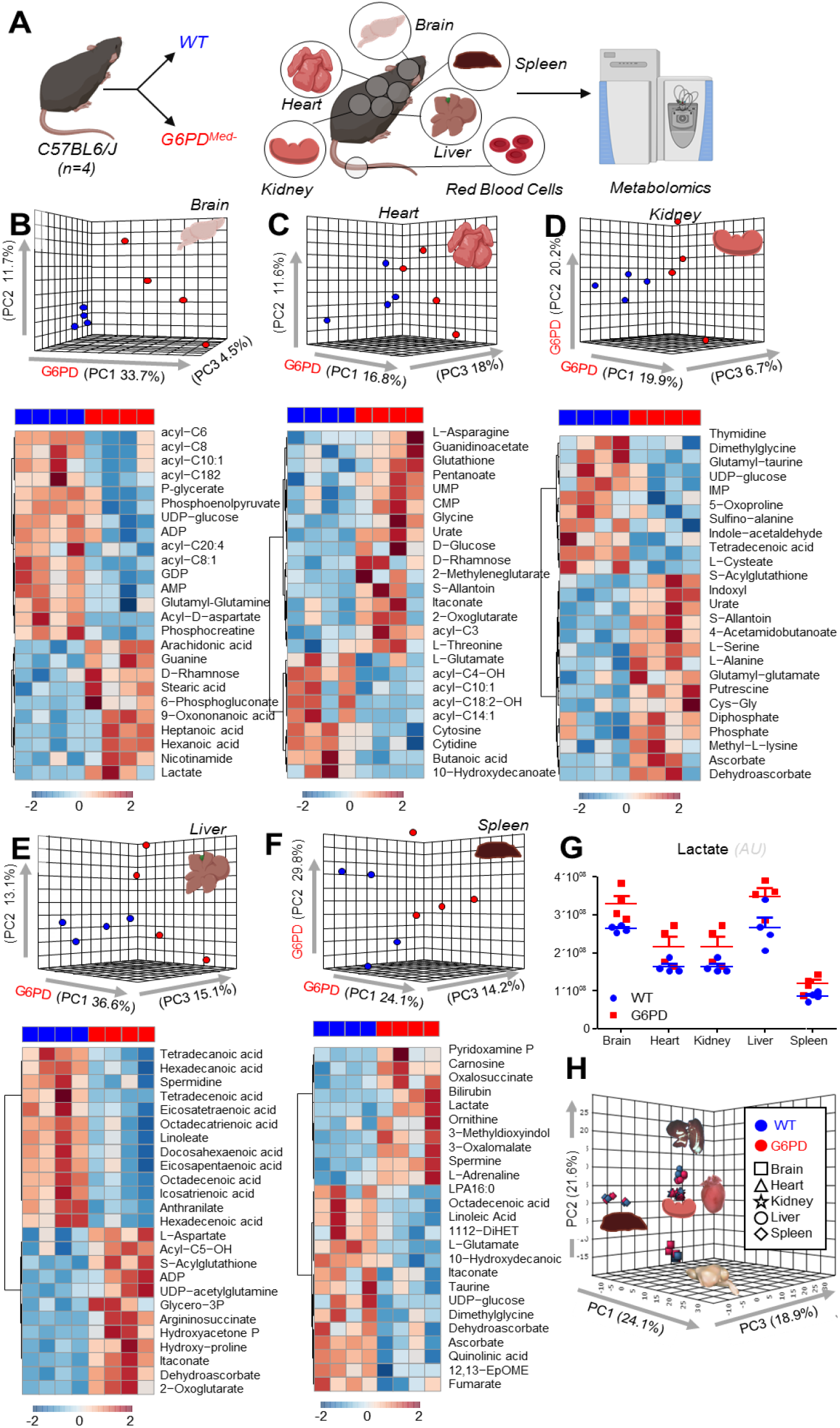
Metabolic analyses of brain, liver, heart, kidney and spleen from WT and G6PD def mice. **(A).** Significant metabolic changes were observed in all the organ tested from transgenic mice, compared to WT controls (**B**-**F**), as highlighted by multivariate principal component and hierarchical clustering analyses (only the top 25 significant metabolites – T-test are shown). The majority of metabolic changes in organs of G6PD subjects related to fatty acid metabolism and acyl-carnitines, followed by amino acid metabolism and glycolysis. Lactate levels were significantly lower in all organs of G6PD deficient mice (**G**), though organ-to-organ metabolic differences overweighed differences between WT and transgenic mice (**H**).

Of note, significant decreases in carnitine-conjugated fatty acids and increases in free fatty acids were noted in all tissues of G6PD_Med-_ mice, most notably in high oxygen-consuming organs, such as the brain (18.4%), heart (11.6%), and liver (20.4%). There was a distinct change in beta-oxidation of unsaturated fatty acids, a hallmark of oxidative metabolism and an NADPH-dependent process(20), suggesting a potential mechanistic linkage between a reduced capacity to sustain NADPH-generation via the PPP in G6PD_Med-_ mice and altered fatty acid metabolism. Similarly, several organs exhibited decreased levels of high-energy phosphate compounds (e.g., ADP, AMP, GDP, phosphocreatine), increased levels of amino acids (metabolized in mitochondria), and increased levels of carboxylic acids (e.g., 2-oxoglutarate and itaconate; **Figure 7B-F**), further suggesting depressed mitochondrial metabolism. Consistent with an apparent depression of oxidative metabolism, all organs also exhibited increased levels of lactate (**Figure 7G**). However, multivariate analyses (**Figure 7H**) indicate that - in the absence of oxidant stress - the metabolic differences among organs are more pronounced than the metabolic differences observed within the same organ in G6PD_Med-_ and wild-type mice.

## Discussion

Herein we describe a murine model of G6PD deficieny that overcomes the limitations of existing models, and has allowed an exploration into metabolomics of both RBCs and solid organs. These data provide novel insight into the effects of an enzymopathy that affects approximatley 5% of all humans. The extent of the decreased activity G6PD_Med-_ mice, as compared to wild-type, approximates what is seen in humans (i.e. 5-10% of normal activity) and, as is the classic finding in humans, pharmacological oxidative stress induces brisk hemolysis in vivo with hemoglobinuria. Metabolic tracing experiments demonstrated a significant decrease in glucose flux through the PPP and increased sensitivity to pharmacological oxidative stress. Although a detailed metabolomics of sold organs in humans with G6PD deficiency has not been reported, the decreased PPP in organs from the G6PD_Med-_ mouse is consistent with known mild decreases in G6PD organs in humans(21-25).

Human G6PD-Med(-) subjects have a mildly shortened RBC circulatory lifespan (i.e., 120 days in G6PD-normal controls, as compared to 100 days in G6PD-Med(-) subjects)(26). In contrast, RBCs from the hMed(-) mice have a normal RBC circulatory lifespan. It can be speculated that the murine RBC lifespan (55 days) may not be long enough for the defects to accumulate, as they do in humans over 120 days. Conversely, in their normal state, mice have higher GSH levels than do humans, which may also mitigate the effects of G6PD defieciency in the absence of external oxidative challenge. Alternatively, other pathways involved in glutathione synthesis (e.g., ascorbate synthesis) exist in mice, but not in humans (27).

To our knowledge, this report contains the first metabolomic analysis of RBC changes after exposure to phenylhydrazine (in either WT of G6PD deficient mice); however, one caveat with this is that the changes could reflect metabolites in newly formed reticulocytes vs alterations in RBCs at the time of treatment. In vitro analysis of diamide treated RBCs does not suffer this issue as new RBC generation is absent. Moreover, detailed metabolomics have been reported following *in vitro* exposure of human G6PD-def RBCs to diamide(28). In particular, it was reported that G6PD-def RBC (but not normal RBCs) responded to diamide though depletion of GSH through oxidation (e.g decreased GSH with increasing GSSG). In response, GSH synthesis was increased as evidenced decreases in GSH precursors (e.g. γ-glutamylcysteine) and generation of byproducts of GSH sythesis (e.g. ophthalmate). ATP was also decreased in G6PD-def RBCs, which was interpreted as reflecting consumption of ATP by GSH synthesis. This resulted in a substantial increase in glycolytic metabolism in G6PD-def RBCs, presumably to compensate for the energy depletion of ATP consumption; however the glycolysis was abnormal in that pyruvate levels did not increase, suggestive of oxidative deactivation of pyruvate kinase. As expected, down stream metabolites of the PPP were decreased in G6PD-def RBCs. The authors of this paper also showed that the increase in glycolysis of G6PD-def RBCs in resposne to diamide correlated with increased AMPK activity that was associated with increased AMP levels due to ATP consumption.

Consistent with the above studies, when we exposed G6PD_Med-_ mouse RBCs to diamide, we observed significant alterations of glutathione metabolism, accompanied by the inability to preserve non-deaminated high-energy purine pools in the face of decreased PPP activity. This is relevant in that RBC purine deamination by oxidant stress-activated AMP deaminase 3 is a hallmark of impaired energy metabolism, ultimately associated with a decreased ability of the RBC to circulate (29) following oxidant insult, and an overall shorter RBC lifespan in mice that are unable to replenish the ATP pool (e.g., mice with altered AMPK activity)(30). Of note, increased purine deamination by AMP deaminase 3 activation phenocopies the protective effect of G6PD deficiency on malaria infection.(31) However, a number of specific metabolic differences were also observed between diamide responses of G6PD_Med-_ mouse RBCs and that reported with human RBCs. Surprisingly, GSH levels did not decrease in mouse G6PD-def RBCs in resposne to diamide treatment; however, GSH levels did decrease in resposne to other oxidative stress (e.g. PHZ). It is worth noting that in the human studies, only a small number of humans were studied, they had a different G6PD mutation (Canton and not Mediterranean) and there may also be methodological differences. Future study will be required to assess the meaning of these respective biologies.

It was recently reported that RBCs from G6PD-deficient human volunteers stored more poorly than G6PD-normal RBCs(32). However, the decrease in post-transfusion recovery was modest (∼6%). Importantly, most subjects in this study had the milder A-form of G6PD deficiency. Only a single subject was of the G6PD-Med(-) type, and unexpectedly, that subject had a post-transfusion recovery overlapping with the median measurment from the control group. The basis for this is unclear, but it indicates that the current findings with the G6PD_Med-_ mouse are not in conflict with the one human G6PD-Med(-) who has been studied. There are conflicting reports regarding the extent to which G6PD activity decreases in RBC storage; it is unclear if these differences were due to differing populations or methodologies (e.g. differences in storage additives such as SAGM vs. AS-3)(33, 34). However, changes in metabolism from the stored RBCs from the G6PD-deficient mice are laregly consistent with larger metabolic studies on RBCs from human donors with the G6PD-Med(-) variant (e.g. dopamine and pyruvate/lactate ratio); although post-transfusion recovereis were not reported for this group(35). Finally, recent findings from the REDS study showed similar findings during storage of units from G6PD deficient donors, although these were largely of the A-variant(36).

Although studies outside the RBC compartment in G6PD-deficient humans are limited, decreased G6PD activity in non-erythroid compartments has been reported, including in leukocytes(24), platelets(25), liver(23), and muscle(21, 22). Similarly to what has been reported in humans, we show mild decreases in PPP activity in multiple organs, consistent with a mild decrease in G6PD activity. The milder deficiency in non-erythroid organs is presumably because, unlike RBCs, they have ongoing synthesis of the destabilized enzyme. More importantly, we provide a comprehensive metabolic description of mulitple organs from G6PD deficient mice, showing a role for altered PPP activation in the cross-regulation of several NADPH-dependent pathways in non-erythroid tissues. Most importantly, the current data highlight a G6PD deficiency-dependent alteration in fatty acid amounts, unsaturation, and metabolism (as gleaned by the levels of free and carnitine-conjugated fatty acids). Notably, fatty acid synthesis (a key anabolic requirement for rapidly proliferating cells), desaturation, and catabolism are NADPH-dependent processes.

The role of NADPH-dependent lipid metabolism is being increasingly appreciated in tumor biology(37). Thus, our observations, combined with the appreciation of the high penetration of G6PD deficiency in human populations, raise the possibility of a previously unappreciated role for G6PD status and biology in solid tumors, analogous to what has been proposed for hematological malignancies(38). In addition, our observations suggest an indirect role for G6PD status on the capacity to metabolize fatty acids in high-oxygen demand organs (e.g., brain, heart, liver), making this model potentially useful for the study of how G6PD deficiency may affect other disorders that have been associated with G6PD deficiency in humans, including kidney disease and diabetes(7), pulmonary arterial hypertension(9), amytrophic lateral sclerosis(8), Huntington Disease(8), Parkinson’s Disease(8), Alzheimer Disease(8), and multiple scerlosis(39).

In summary, we report a new model of G6PD deficiency in mice using a humanized enzyme of the Med-variant, and describe novel metabolic findings with regards to normal RBC biology, oxidant stress responses, and systemic alterations in peripheral organs. This new model promises to be of high utility in ongoing studies in a wide variety of biologies and pathologies in which G6PD deficiency has been implicated (e.g. aging(40), pulomnary hypertension, neurological disorders(8), exercise physiology, anti-malarial toxicology(41), mechanisms of genetic resistance to malarial infection, etc.).

## Material and Methods

### Mice

The Con-hMed(-) were generated, as described in detail in the results section, using Bruce4 ES cells. Con-hMed(-) mice were bred to mice expressing CRE under a CMV promoter (Jackson mouse (B6.C-Tg(CMV-cre)1Cgn/J stock # 006054)) resulting in G6PD_Med-_ mice, which were maintained through backcrossing to C57BL/6J mice. Female heterozygous G6PD_Med-_ mice were bred with wild-type C57BL/6J mice such that all wild-type and G6PD_Med-_ mice were littermate controls from the same breeding colony. Ubi-GFP (UbiC-GFP (C57BL/6-Tg(UBC-GFP)30Scha/J stock # 004353)) and C57BL/6J mice were purchased from Jackson Labs and bred as described. All experiments were carried out under an approved IACUC protocol in the BloodworksNW vivarium and all mice were used from 2-6 months of age – for any given experiment mouse ages were matched.

### RNA isolation/RT-PCR

Bone marrow was harvested from mice, and mature RBCs lysed with RBC lysis buffer. Remaining cells were resuspended in Trizol (ThermoFisher), and RNA extraction was performed per the manufacturer’s instructions. RNA was converted to cDNA using the iScript gDNA clear kit (BioRad). Real-time qPCR was performed on a QuantStudio 6 Flex (Applied Biosystems), using the following Taqman primers and probes: ActB (Mm00607939_s1), Pol2Ra (Mm00839502_m1), murine G6PDx (Mm04260097_m1; spans exons 1-2), murine G6PDx (Mm00656735_g1; spans exons 12-13), human G6PD (Hs00959072_g1; spans exons 3-4), and human G6PD (Hs00959073; spans exons 4-5). Controls lacking reverse-transcriptase were run for every RNA preparation to control for the presence of genomic DNA; none of these samples amplified (data not shown). Data were analyzed with the QuantStudio 6 software using relative quantitation (ΔΔCt), with ActB as the normalization factor and B6 as the control strain.

### RBC lysis and hemoglobin depletion

RBC ghosts (membranes) were prepared by hypotonic lysis, followed by depletion of hemoglobin. Briefly, RBCs were washed 3 times PBS, followed by transfer of one part washed RBCs into three parts water, followed by end over end rotation for 5 min at room temperature to lyse the RBCs. Lysed RBCs were then mixed 1:1 with Hemoglobind (Biotech Support Group), followed by end over end rotation for 10 min at room temperature. Hemoglobind and bound hemoglobin were pelleted by centrifugation, and supernatants subjected to an additional hemoglobin depletion with hemoglobind. Supernatants were used for western blotting (below).

### Western blotting

Hemoglobin-depleted supernatants were electrophoresed on 4-12% NuPAGE Bis-Tris gels under reducing conditions, and transferred to PVDF membranes. As a control, recombinant human G6PD protein (Abcam, cat#NP0007) was also included. Membranes were blocked in 2.5% milk/2.5% BSA in TBST, and probed with rabbit anti-human/mouse G6PD (Abcam, cat# EPR20688) or rabbit anti-Actin (Cell Signaling Technology, cat#4970), and imaged by standard chemiluminescence techniques on an ImageQuat-800 (GE).

### G6PD Activity Assay

For the quantitative determination of G6PD activity, the Glucose-6-Phosphate Dehydrogenase Reagent Set was used (Pointe Scientific, cat# G7583180). Following the manufacturer’s protocol, 10ul of whole non-washed RBCs were assayed using a heated cuvette spectrophotometer (Nanophotometer C40, Implen); Hb(g/dL) was measured using ∼70ul whole, unwashed RBCs on an ABLX90 (Radiometer). Results are presented as G6PD activity (U/g Hb).

### Statistics for RT-PCR

Analysis using two-way ANOVA with Sidak post-hoc testing revealed no significant differences in mRNA levels between WT and G6PD^Med-^ mice for either Pol2RA or Mm04260097_m1.

### Blood Storage and Transfusion Studies

Blood storage, transfusion, and determination of post-transfusion recovery were carried out as described in detail in previous studies(17, 42).

### RBC lifespan determination and oxidant stress challenge

RBC lifespan determination was carried out by the biotinylation method(14), and as described in detail in previous studies(43). Phenylhydrazine was given to mice through 3 intraperitoneal injections of 0.01 mg/g administered 12 hours apart.

### Metabolomics

#### Sample processing and metabolite extraction

Fifty µL of frozen RBC aliquots or 10 mg of snap frozen organ tissues were extracted in 450 ul or 1 ml, respectively, of ice cold methanol:acetonitrile:water (5:3:2 *v/v*). Samples were agitated at 4°C for 30 min followed by centrifugation at 10,000 x *g* for 10 min at 4°C, as described (44). Protein pellets were discarded, and supernatants were stored at -80°C prior to metabolomic analysis.

#### Ultra-High-Pressure Liquid Chromatography-Mass Spectrometry (MS) metabolomics

Samples were randomized and 10 μl aliquots of extracts were injected using a UHPLC system (Vanquish, Thermo, San Jose, CA, USA) and run on a Kinetex C18 column (150 × 2.1 mm, 1.7 μm – Phenomenex, Torrance, CA, USA) at 250 μl/min (isocratic: 5% Optima acetonitrile, 95% Optima H2O, 0.1% formic acid)(45) and 400 μl/min (5 or 17 min gradient 5-95% B; A:water +0.1% formic acid, B: acetonitrile + 0.1% formic acid)(46, 47). Formic acid in mobile phases was replaced by 1 mM ammonium acetate for negative mode runs. The UHPLC system was coupled online with a Q Exactive mass spectrometer (Thermo, Bremen, Germany), scanning in Full MS mode (2 μscans) at 70,000 resolution in the 60-900 m/z range operated in either polarity mode. Eluates were subjected to electrospray ionization (ESI) in positive and negative ion modes (separate runs) with 4 kV spray voltage, 15 sheath gas, and 5 auxiliary gas. Separate Top15 ddMS2 runs were performed on tech mixes composed of 10 μl of each extract from each one of the samples in the batch. Chromatographic and MS technical stability were assessed by determining CVs <10% for metabolites in mixed controls run every 5 injections in the queue. MS1 and data-dependent MS2 acquisition(48), data analysis, and elaboration were performed, as described(46, 47). Graphs and statistical analyses (either t-test or repeated measures or two-way ANOVA) were generated with GraphPad Prism 8.1.2 (GraphPad Software, Inc, La Jolla, CA), GENE E (Broad Institute, Cambridge, MA, USA), and MetaboAnalyst 4.0(49).

## Author Contributions

ADA, SLS, EAH, ROF, MK, TT and JCZ concieved of the studiess, designed experiments and interpreted data. HLH, AMH, BB, MJW, and XF carried out experiments, generated data and interperted data. All authors were actively involved in the writing of the manuscript and the interpretation of data.

## Acknowledgements

This research was supported by funds from the RM1GM131968 (ADA) from the National Institute of General and Medical Sciences, and R01HL146442 (ADA), R01HL149714 (ADA), R01HL148151 (SLS, ADA, JCZ), R21HL150032 (ADA), from the National Heart, Lung, and Blood Institute.

