## Supplementary Figure 1 for "Hematologic and systemic metabolic alterations due to Mediterranean type II G6PD deficiency in mice"

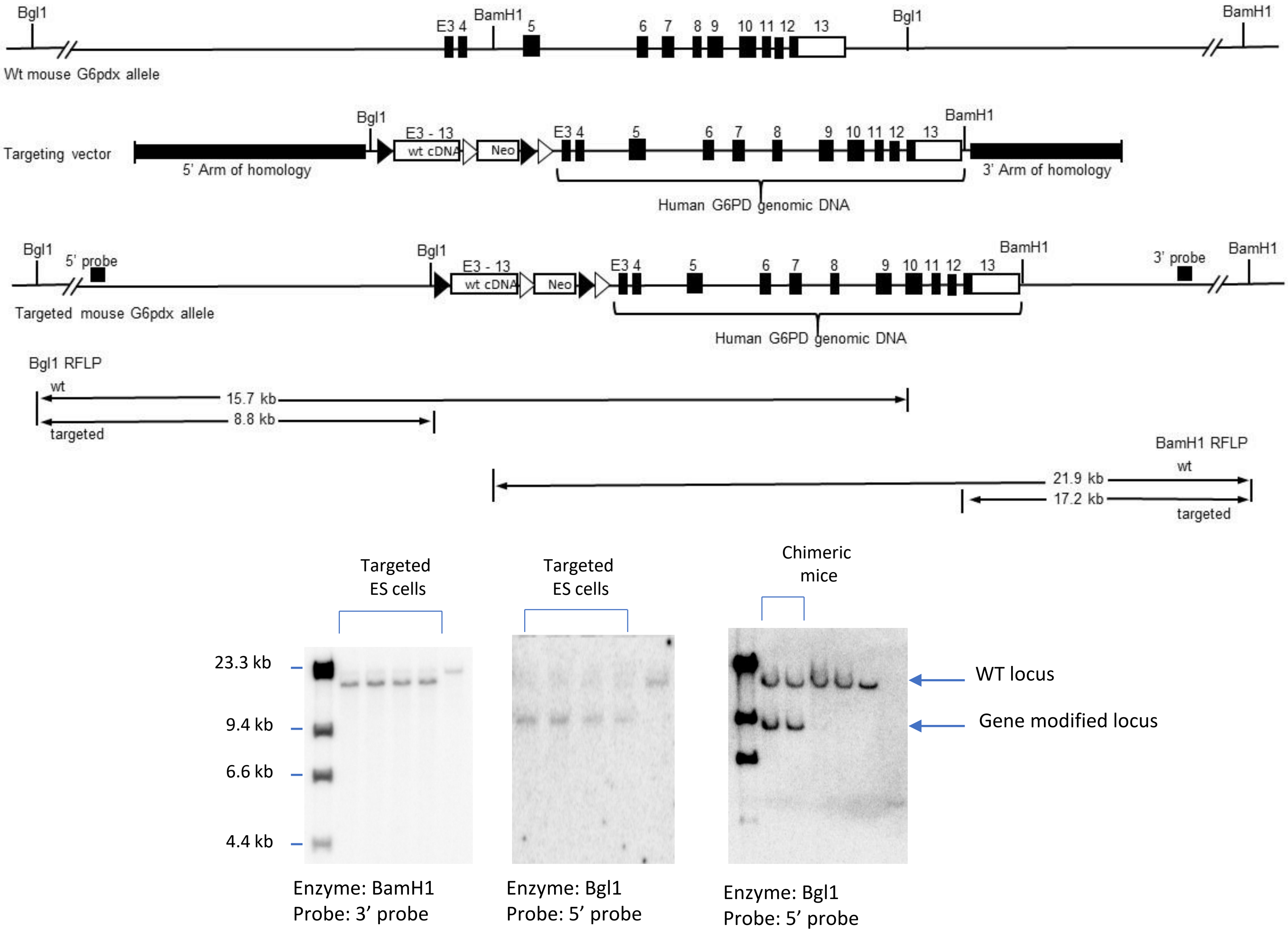


**Supplemental Figure 1.** The schematic shows restriction sites in wild-type (Wt) and gene modified loci. Southern blot analysis of ES cells and the final mice demonstrate homologous recombination and the absence of any random iintegration.
