## Supplementary Figure 2 for "Hematologic and systemic metabolic alterations due to Mediterranean type II G6PD deficiency in mice"

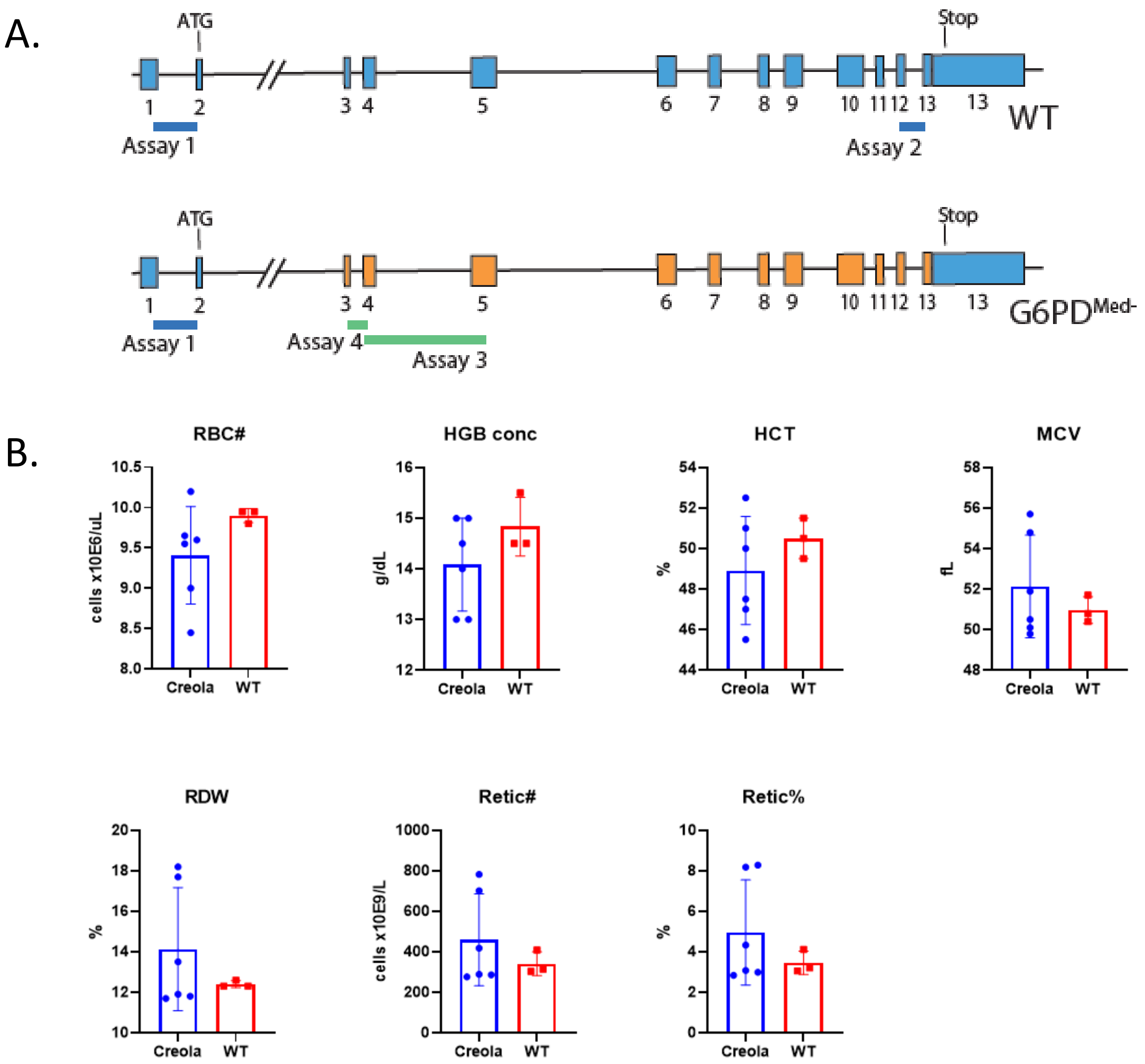


**Supplemental Figure 2. (A) Schematic** of wild-type (WT) and G6PD$_{Med-}$ Genes indicating the location of allele specific probes. **(B)** Values of indicated blood parameters. All bar graphs represent means with SD, no significant differences were observed by Welch's t-test (assumption of Gaussian distribution and unequal SDs)
