## Supplementary figures and images for "Hematologic and systemic metabolic alterations due to Mediterranean type II G6PD deficiency in mice"

### Supplementary Figure 3

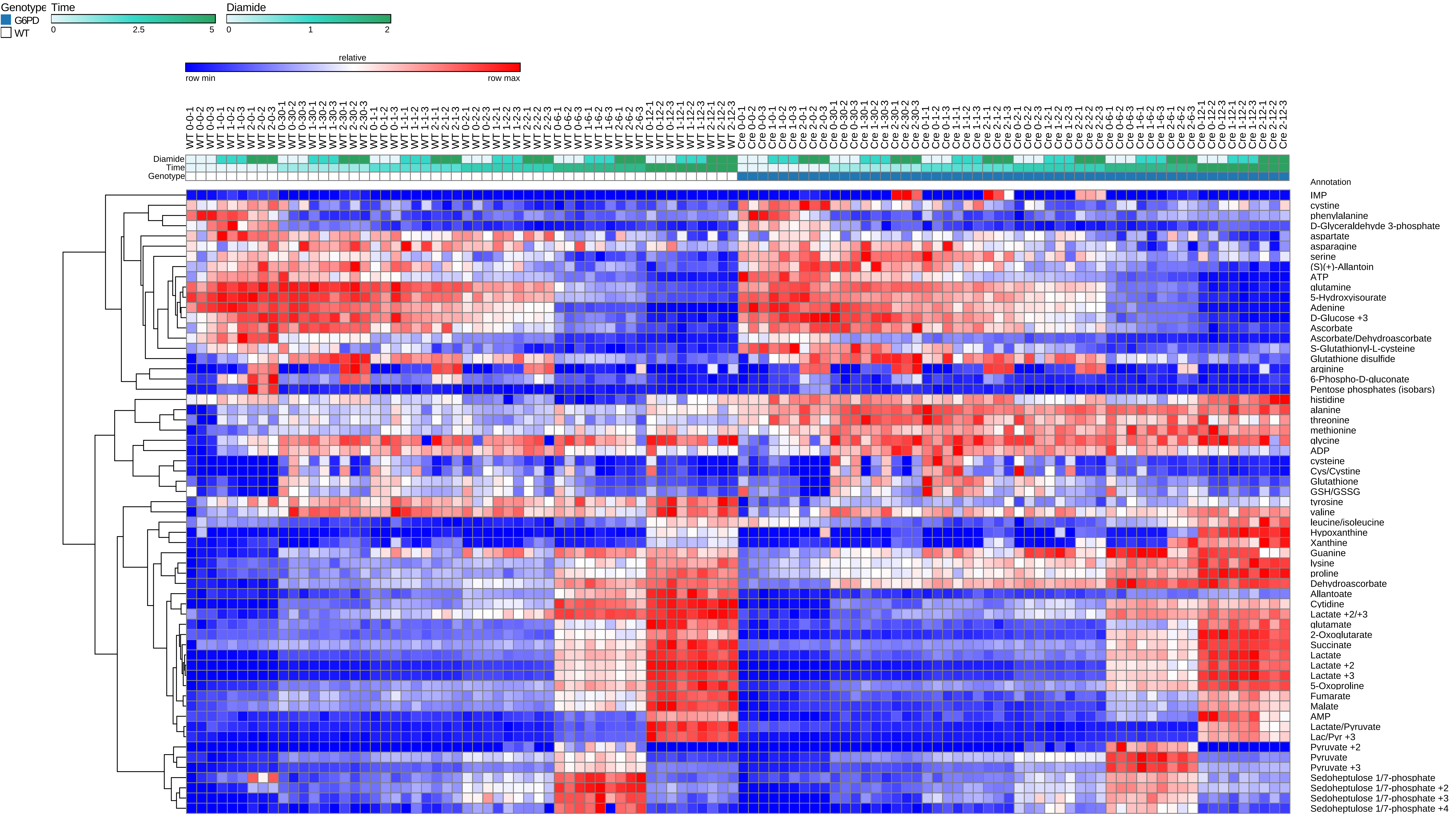

### Supplementary Figure 4

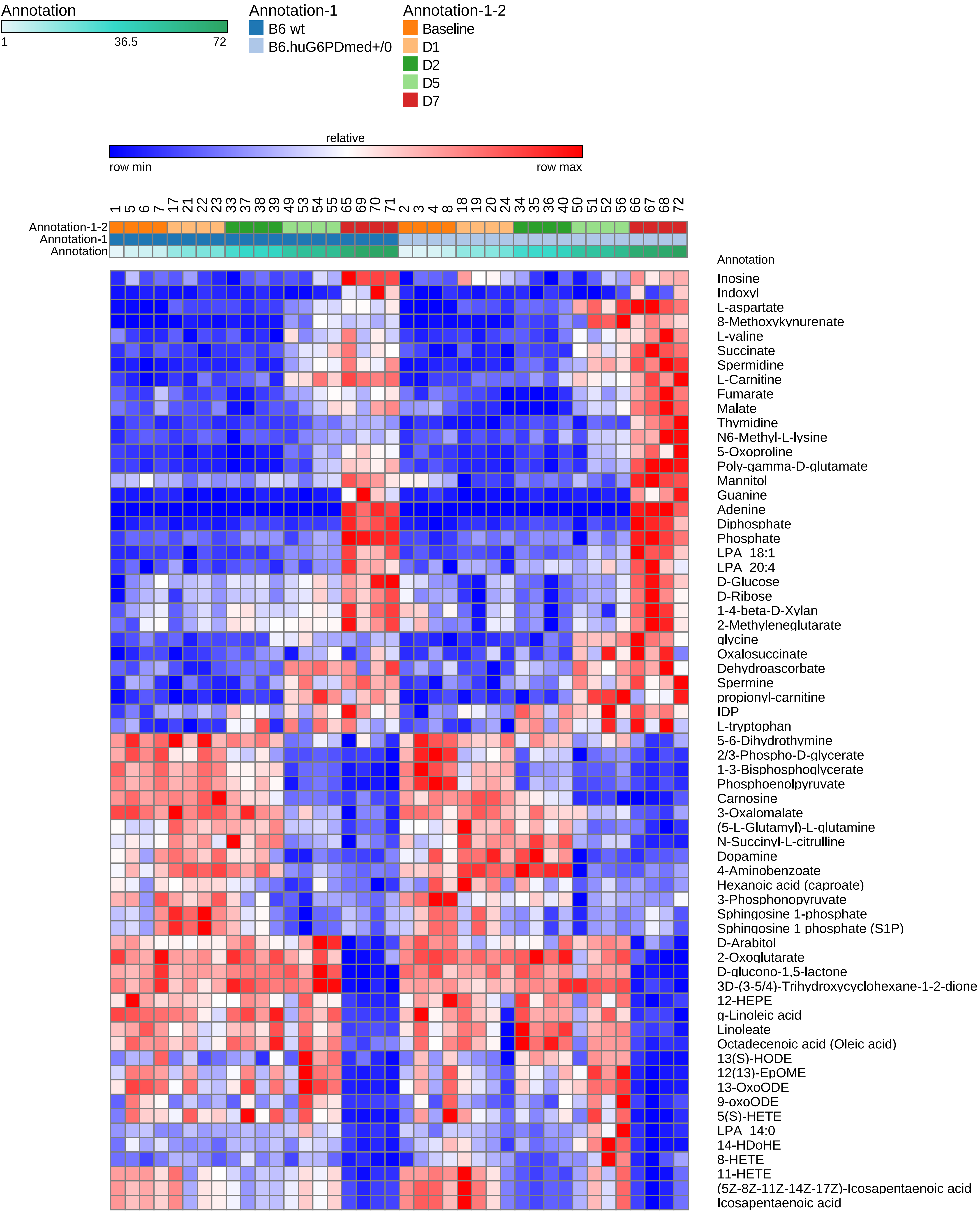

### Supplementary Figure 5

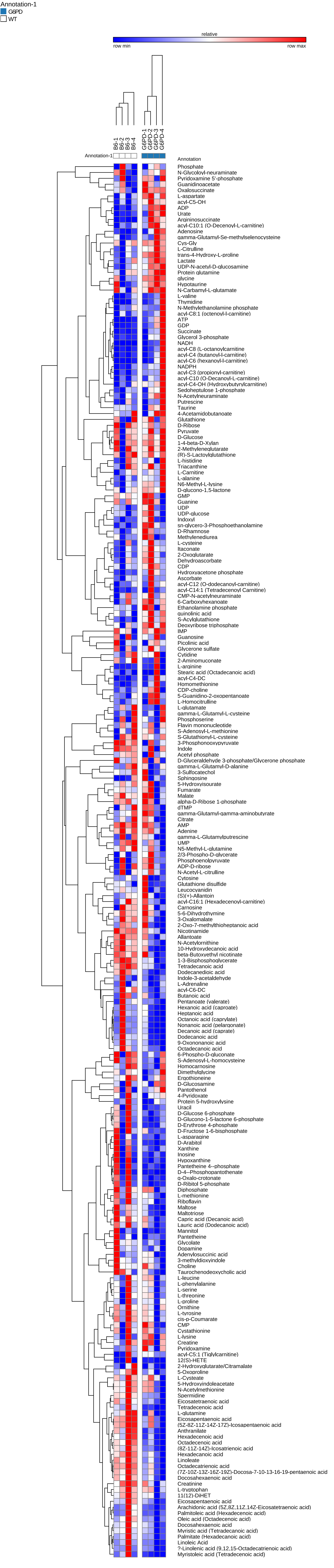

### Supplementary Figure 6

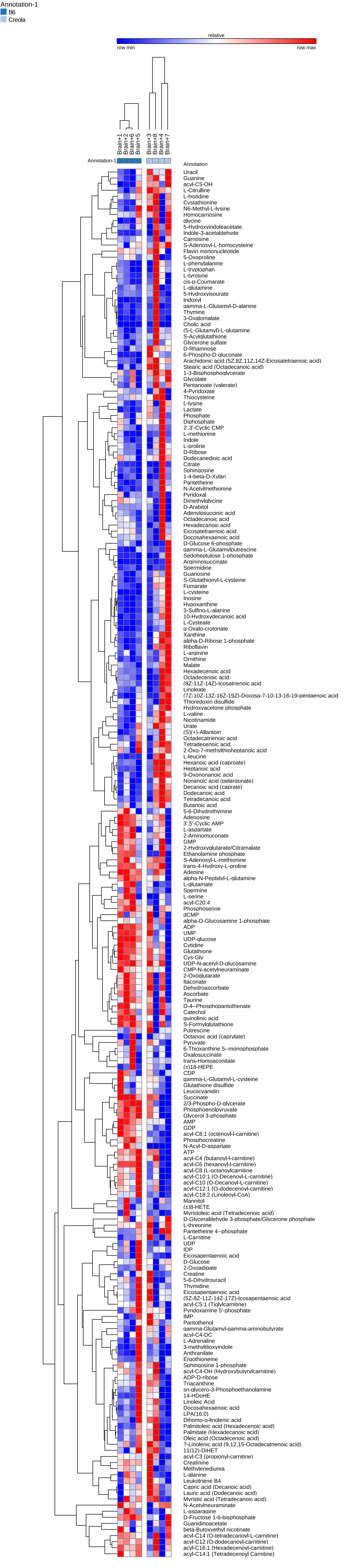

### Supplementary Figure 7

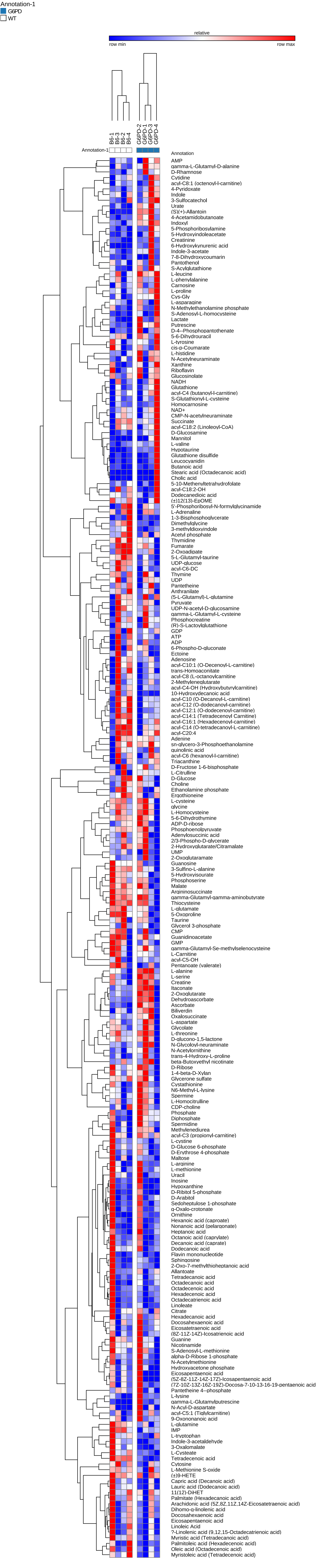

### Supplementary Figure 8

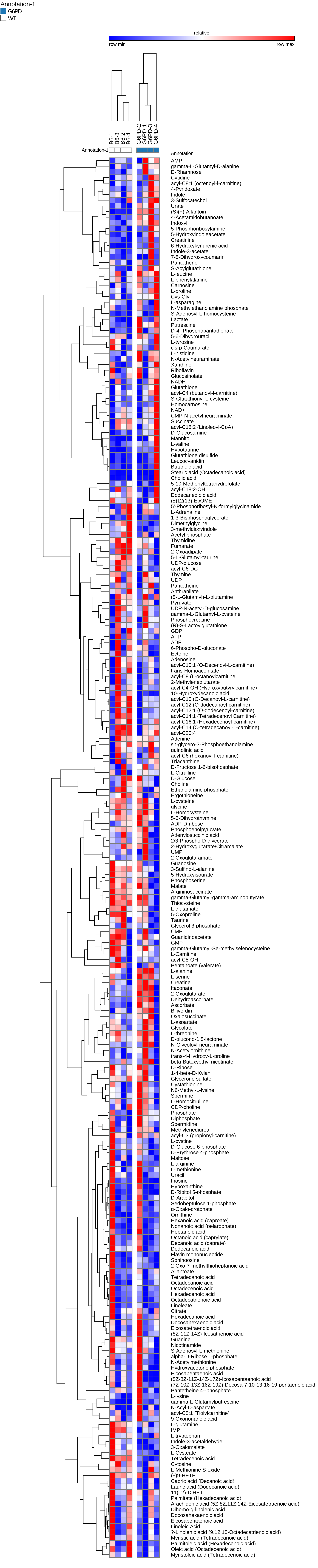

### Supplementary Figure 9

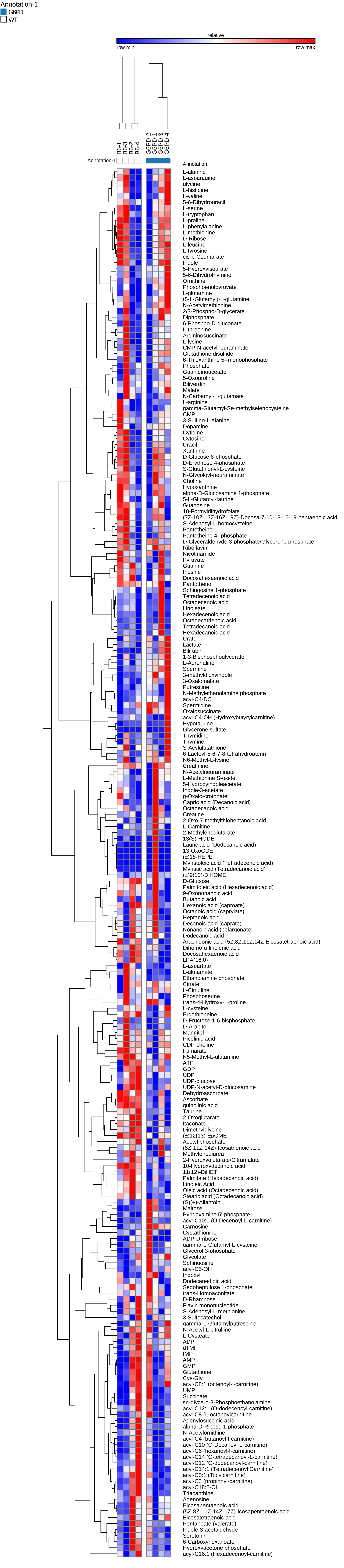

### Supplementary Figure 10

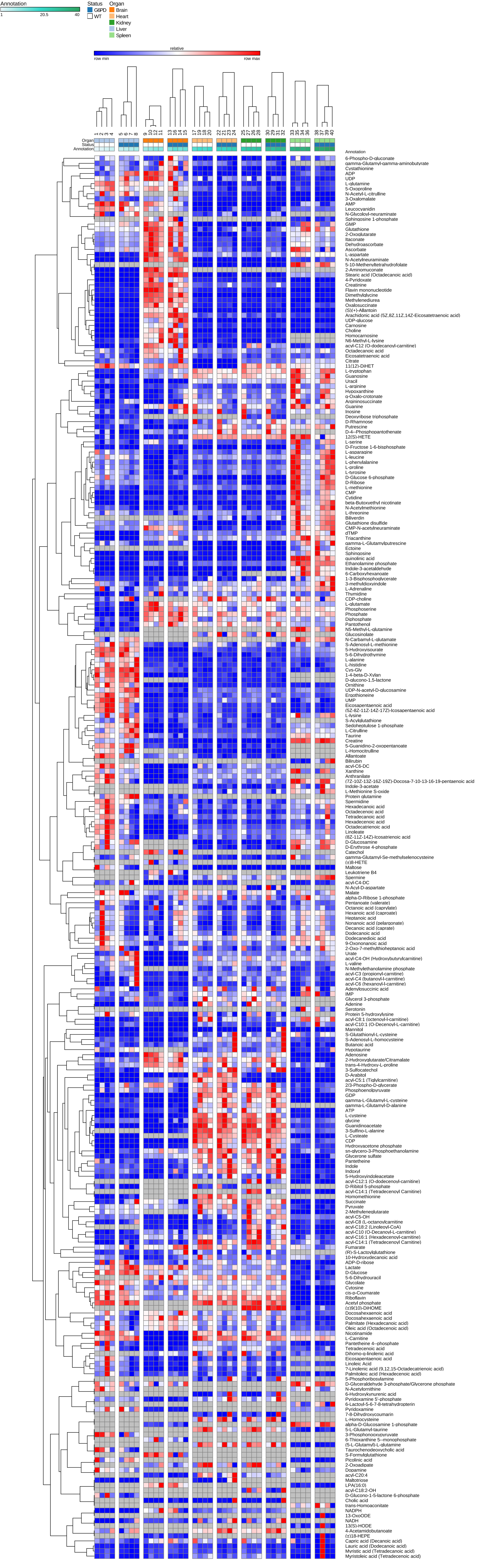
